# Recursive exploration of metabolic yield space

**DOI:** 10.64898/2026.05.28.728453

**Authors:** Wannes Mores, Satyajeet Bhonsale, Stylianos Floros, Filip Logist, Jan Van Impe

**Author notes:** Corresponding author, *Email address:* (Jan Van Impe).

## Abstract

Genome-scale metabolic network reconstructions contain extremely detailed and valuable information regarding cellular metabolism. For many applications such as finding genetic engineering targets and reduced kinetic model construction, metabolic network analysis techniques exist. Yield spaces based on the extreme rays of solution cones related to the metabolic network are frequently constructed for these types of analyses. However, for genome-scale networks, full enumeration of these extreme rays is not computationally feasible. In this work, a novel direct generation method for yield spaces is presented. This allows the application of many metabolic network analysis techniques to even the most recent genome-scale metabolic networks. Inspired by principles from multi-objective optimization algorithms, the proposed method performs highly efficient recursive exploration but specifically adapted to the mathematical properties of yield spaces. Two case studies showcase both the efficiency of the method and its applicability for analysis of genome-scale metabolic networks.

## 1. Introduction

In computational biology, metabolic networks are frequently used as a tool to predict organism behaviour and analysing its metabolic capabilities *in-silico*. These networks are defined by the stoichiometric matrix **S** ∈ ℝ^*m×r*^, which maps the all *m* metabolites to all reactions *r* they take part in. By applying the pseudo-steady-state assumption, we get the following linear system that lies at the basis of constraint-based modelling:

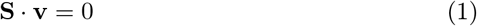

Throughout recent years, metabolic networks have become more comprehensive. While these genome-scale metabolic networks (GEMs) contain highly relevant information, they are also high in complexity, making their integration challenging in many applications. Therefore, metabolic network analysis tools are vital to efficiently extract and use the relevant information on cellular metabolism present in the network.

### 1.1 Metabolic network analysis

One of the most powerful analysis tools for metabolic networks is through its extreme rays of the flux cone. Depending how reversible reactions are dealt with, different types of rays can be generated. The most commonly used rays are elementary flux modes (EFM) [21], elementary flux vectors [24] and extreme pathways (EP) [19]. An overview of the differences in their generation, their properties and uses can be found in Llaneras & Picó [7]. A common problem when applying the original generation methods for extreme rays is the combinatorial explosion of intermediate candidates. For GEMs, this leads to such a large set of candidates that it cannot be stored in memory. Alternatively, partial enumeration of the rays for GEMs is possible [14, 17], but cannot guarantee full coverage of metabolic capabilities of the original network.

For larger networks, the total number of EFMs or EPs is vast, requiring appropriate selection strategies. Yield analysis [23] allows for a large reduction by only considering the rays that lie on the edge of the yield space. These yield spaces are defined by evaluating the yield of each ray under consideration for a list of products in relation to a chosen substrate. By selecting only the points on the convex hull, the entire yield space is still generatable through non-negative combination of the remaining rays. Further reduction is then possible by considering minimal sets that keep a chosen percentage of the volume of the yield space, often 99%.

### 1.2. Applications of metabolic network analysis

Determining appropriate extreme rays for metabolic networks has been proven very useful in many applications. For metabolic engineering, EFMs have often been used to define engineering targets. One example for this is their use in defining minimal cut sets (MCS), which contain combinations of gene or reaction deletions that block certain unwanted metabolic behaviours [4]. More recently, an alternative route to generating MCS was found, where individual EFMs of a dual system of the metabolic network directly leads to one MCS, removing the need for full EFM sets [6, 11].

Another application of extreme rays and yield spaces is the creation of bioprocess models. The bioprocess is represented by a very small set of rays, which is selected either through solving many estimation problems [10], determining a set of points in yield space which capture the measured yields [23], or a combination of both [13]. For a quality reduced model, the original set of EFMs or EPs has to cover enough of its metabolic capabilities, therefore often relying on full EFM or EP sets, which is why it has not been applied yet to genome-scale metabolic networks.

### 1.3. Direct generation of yield spaces

As discussed, yield spaces have mostly been constructed through EFM analysis. This would limit their use to medium-scale networks as EFM enumeration is challenging for GEMs. However, recently, some approaches have been created to directly generate the yield space by solving the associated linear fractional program (LFP):

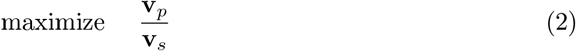

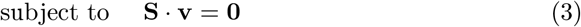

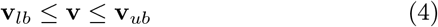

Where **v**_*p*_, **v**_*s*_, **v**_*lb*_ and **v**_*ub*_ are the product flux, substrate flux, lower bounds on the flux vector, upper bounds on the flux vector respectively. Luo et al. [8] transform the problem to a series of LPs that is proven to converge to the maximal yield. These LPs are very fast and often require only two solves, making it a very attractive approach. Alternatively, Klamt et al. [5] and the StrainDesign package [20] transform the LFP to a single LP using the Charnes-Cooper transformation, which significantly enlarges the constraint matrices.

All current approaches for direct generation of yield spaces are aimed at generating two-dimensional spaces. In some cases, a third dimension is added by sampling in the two-dimensional yield space and finding the maximum and minimum yield for each point in the third dimension [20]. These kinds of approaches are very inefficient and scale poorly to higher-dimensional spaces. In multi-objective optimization, high dimensional feasible spaces are already explored efficiently towards determining the Pareto boundary. By introducing a division scheme of the CHIM hyperplane, the space can be explored recursively, allowing for specific regions to receive higher attention as they are deemed more important or informative [15, 3]. Current yield space generation approaches would greatly benefit from a similar approach, allowing the use of more informative, high-dimensional yield spaces for metabolic engineering and construction of dynamic models.

### 1.4. Scope of this work

High-dimensional yield spaces contain the necessary information for reduced model construction and help predict and understand the consequences of metabolic engineering target. Therefore, the scope of this paper is making their generation possible for GEMs. By taking inspiration from multi-objective scalarization approaches, a novel algorithm is defined based on recursive exploration of the high-dimensional yield space of interest. However, significant adaptations have to be made as not only the Pareto optimal points are of interest in the case of yield spaces. A novel recursive division scheme is therefore implemented, together with a termination criterion specific for convex polytopes, which includes yield spaces under standard FBA assumptions

## 2. Methods

In this section, the different building blocks needed for the direct generation of high-dimensional yield spaces are first defined. Following this, an overview of the novel algorithm is given. Lastly, two case studies are described which are aimed at showcasing the efficiency of the algorithm in a benchmark and the applicability to genome-scale metabolic networks respectively.

### 2.1. Direct generation of high-dimensional yield spaces

#### 2.1.1. LP reformulation

To move to higher dimensional yield space generation, the LP formulation is adapted such that multiple yields are optimized simultaneously. The formulation from Luo et al. [8] is used as a basis for the reformulation as it allows very fast optimization of yield. For a single yield based on a selected product *v*_*p*_ and a substrate *v*_*s*_, this LP is defined as:

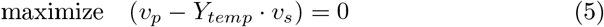

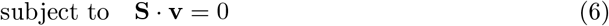

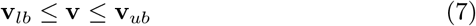

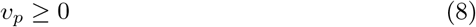

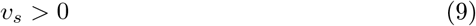

where *Y*_*temp*_ is an initial guess of the yield, updated after an initial solve of the LP using the optimized fluxes. The LP is rerun until the term to be maximized is zero, indicating that *Y*_*temp*_ has converged and the maximal (or minimal) yield *y*^*max*^ = *Y*_*temp*_. A more detailed description can be found in Luo et al. [8]. This LP will be applied later on to find the specific minimum and maximum values for individual yields within CHIM construction.

Taking inspiration from the Normal Boundary Intersection or NBI [1] reformulation of optimization problems, a new variable *e* is introduced which is tied to distance along a vector in the yield space ***λ***. In NBI, the Convex Hull of Individual Minima (CHIM) serves as the reference for creating the vectors, which in the context of multi-objective optimization is the *n −* 1 dimensional simplex formed by the individual minimizers of all *n* objective functions. For the proposed method in this work, a similar approach is adopted, but the CHIM definition is slightly adapted (See Section 2.1.2). The vector ***λ*** in the proposed method is defined by the direction from the nadir point to a given point on the CHIM, a visual example of such a vector can be seen in Figure 1. Additionally, the constraint in Equation 11 is added, which ensures that the generated yield point will lie on this vector. A full explanation of the reformulation steps can be found in the Supplementary Material, with the resulting LP defined as follows:

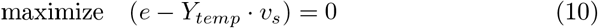

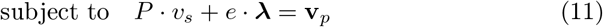

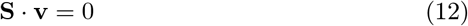

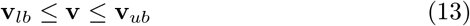

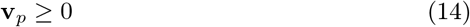

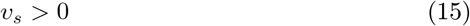

**Figure 1.**
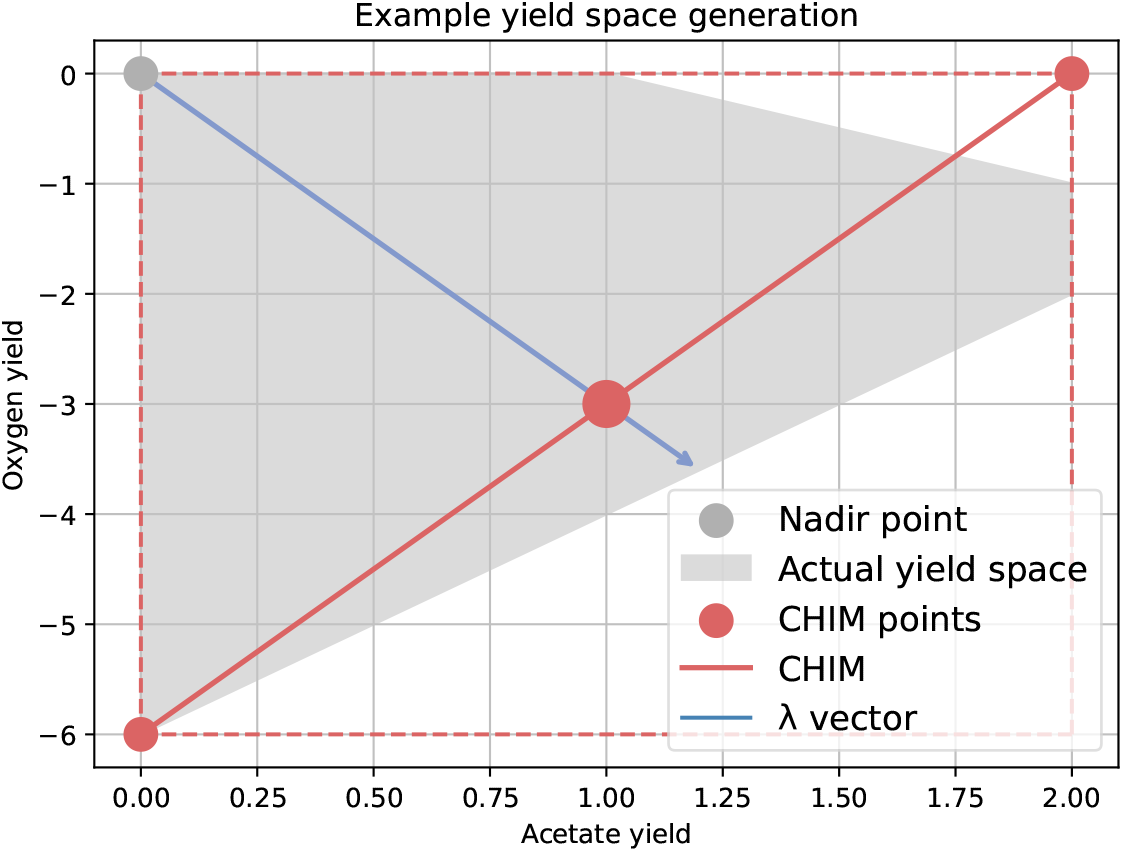
Example construction of the adapted CHIM. In this example, the adaptation to CHIM construction is visible, taking the point (2.0, 0.0) as an edge of the CHIM instead of (2.0, −2.0). This allows for exploration of the full yield space, as opposed to just focusing on the Pareto front.

#### 2.1.2. CHIM creation

Since the goal of the yield space exploration algorithm is to cover the entire space and not only the Pareto optimal points, the CHIM construction from NBI has to be adapted. Each yield of interest is used as a minimization and maximization objective for the LP defined by Equations (5-9), resulting in yield vectors **y**^*min,i*^ and **y**^*max,i*^. The nadir point is then 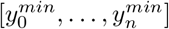 corresponding to the point consisting of all individual minimal values of each yield. In this case, *y*^*min*^ is the *i*th yield in yield vector **y**^*min,i*^. If we consider a certain number of yields *n*, the adapted CHIM is then an *n −* 1 dimensional simplex. Each point is first filled with all the values of the nadir point, with each time the one yield substituted with the optimal value for that yield. For example, the point corresponding to the *i*th yield would be 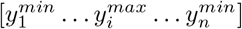 .

By using this adapted CHIM creation, the entire feasible space can be covered by drawing vectors originating from the nadir point and pointing towards each point of the CHIM separately. A 2-dimensional example of this can be seen in Figure 1.

#### 2.1.3. Simplex subdivision scheme

With the reformulated LP and definition of the adapted CHIM, yield points can now be generated once a vector ***λ*** is chosen. A recursive scheme is used to explore different areas of the yield space by applying the Divide & Conquer principle on the CHIM [15, 3]. Barycentric subdivision is used, which allows the CHIM, a simplex, to be divided into smaller simplices of the same *n* dimensions. A visual example of barycentric subdivision for a 2-dimensional simplex can be seen in Figure 2. Alternatively, a hypercube division scheme could be adopted, which would divide the CHIM recursively into hypercubes as described in Hashem et al. [3].

**Figure 2.**
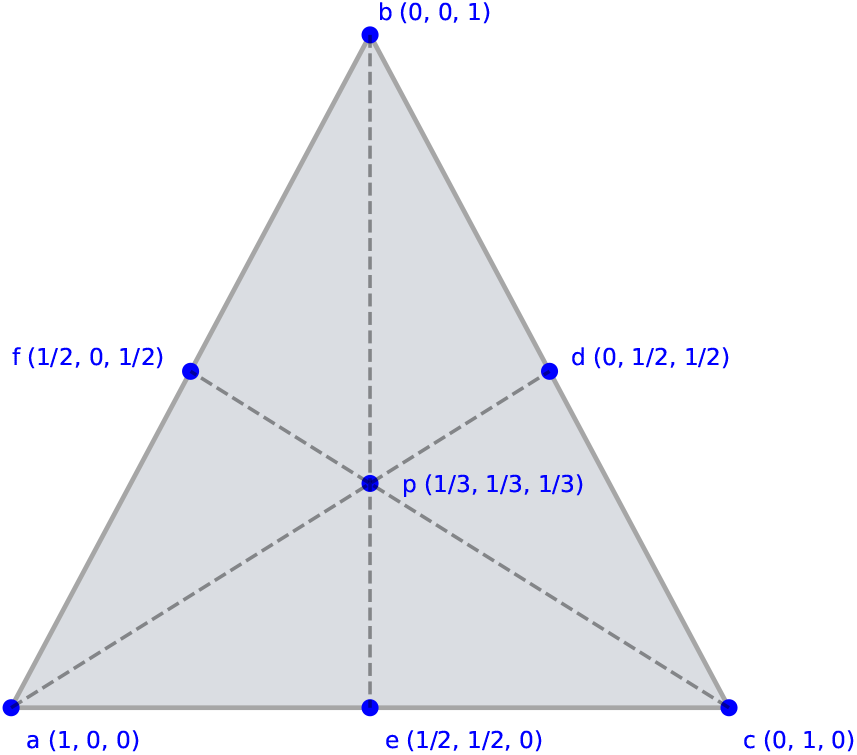
Visual example of barycentric subdivision for a 2-dimensional simplex. The main simplex (a b c) is split into 6 smaller simplices, for example simplex (a e p).

#### 2.1.4. Stopping criterion

To complete the Divide & Conquer principle, a stopping criterion has to be defined, stopping the recursive division in areas of the yield space where sufficient information is retrieved. The main assumption made for the definition of the stopping criterion is that all product yields are bounded, which ensures the yield space is a convex polytope [5]. This assumption relies on good selection of yield spaces, not allowing product formation without the reference substrate not being consumed. If that were possible, it is very likely some other substrate is used and should receive its own yield space(cf. glucose and xylose yield spaces in Song & Ramkrishna [23]). With proper modification of the medium of the model, these yield spaces can be explored separately.

Defining a stopping criterion for convex polytopes can be done geometrically, halting exploration of space when the boundary is found. By using the definition of faces of convex sets [18], it is sufficient to check whether some strictly interior point of a simplex Δ (a convex subset) is on the boundary to prove that the entire simplex is contained within a face of the convex set. From the vertices **x** ∈ Δ, projections **x**^*^ can be calculated from their location on the CHIM to the boundary of the yield space are obtained by solving the LP defined by Equations (10-15). This results in a projected simplex Δ^*^. The same LP is solved for the midpoint of the simplex on the CHIM **p**, resulting in **p**^*^. What remains to fulfill the definition of a face is to check whether **p**^*^ lies on the interior of the projected vertices **x**^*^.

A simple way to verify whether a point lies on the interior of a set of *n* vertices of a simplex is by checking whether the *n*th hypervolume of the convex set defined by these points is zero, which lies at the basis of the stopping criterion. Algorithm 1 provides a definition of the criterion, using projected vertices 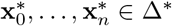 and *ϵ*, a tolerance chosen by the user which defines at which point the volume of the convex hull of points is considered zero.

##### Algorithm 1 Geometric stopping criterion

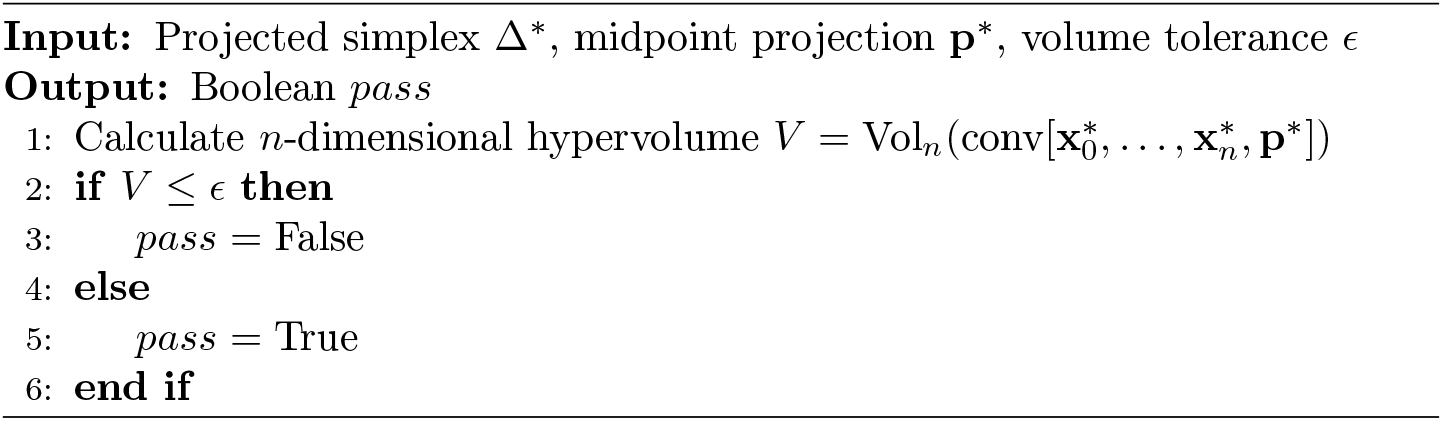

### 2.2. Algorithm description

Combining the steps defined in the previous sections yields the total algorithm for the exploration of higher dimensional yield spaces (Alg. 2). To start, each product of interest defined in **v**_*p*_ is evaluated in terms of its maximal possible value and its minimal possible value. With this information, the initial CHIM is constructed and will serve as the first simplex. Afterwards, the main exploration loop starts, which consists of two parts. The first part will evaluate the current list of simplices, checking whether it could reveal more information regarding the boundary of the yield space by evaluating the geometric stopping criterion. If it could, the simplex is divided using the barycentric subdivision scheme (Section 2.1.3). The second part of the main loop will then evaluate all vertices of the new simplices, adding them to the list of yield points if not already present. When the main loop runs out of simplices, the algorithm stops and returns the final set of yield points *𝒴*.

#### Algorithm 2 Recursive exploration of metabolic yield space

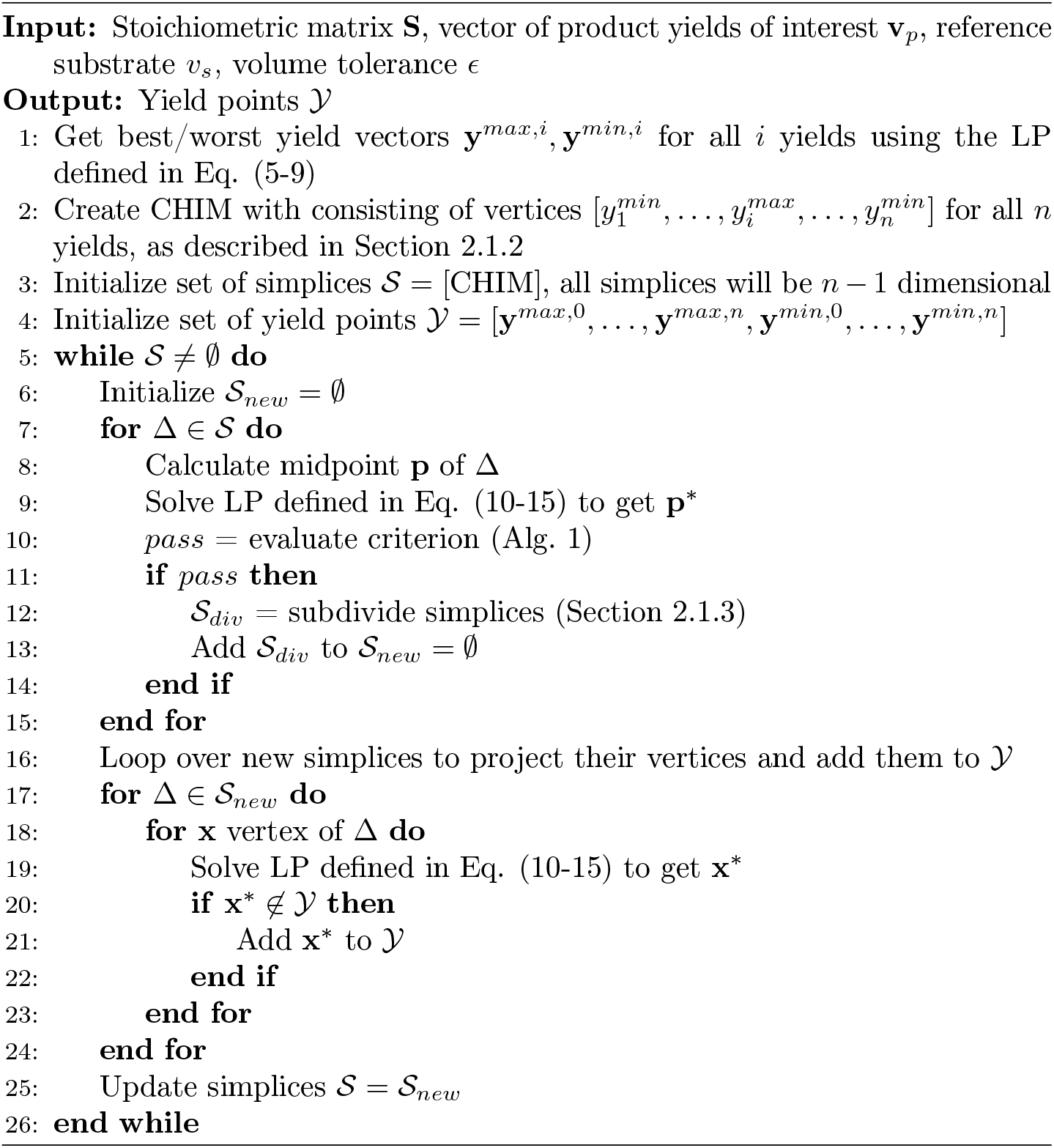

A visual example of recursive algorithm can be seen in Figure 3. A 2D yield space for the e_coli_core model [16] is explored, based on the acetate and oxygen yield where glucose is the reference substrate. The progress of the recursive algorithm is shown at different depth levels. Figures 3b and 3c show the passing and failing of the geometric stopping criterion respectively, where in Figure 3c the 2D simplex has no area which indicates a flat edge of the yield space between 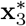 and 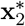 . Therefore, the **x**_3_ *−* **x**_2_ region on the CHIM is not further explored afterwards. Figure 3d then shows the final yield space generated.

**Figure 3.**
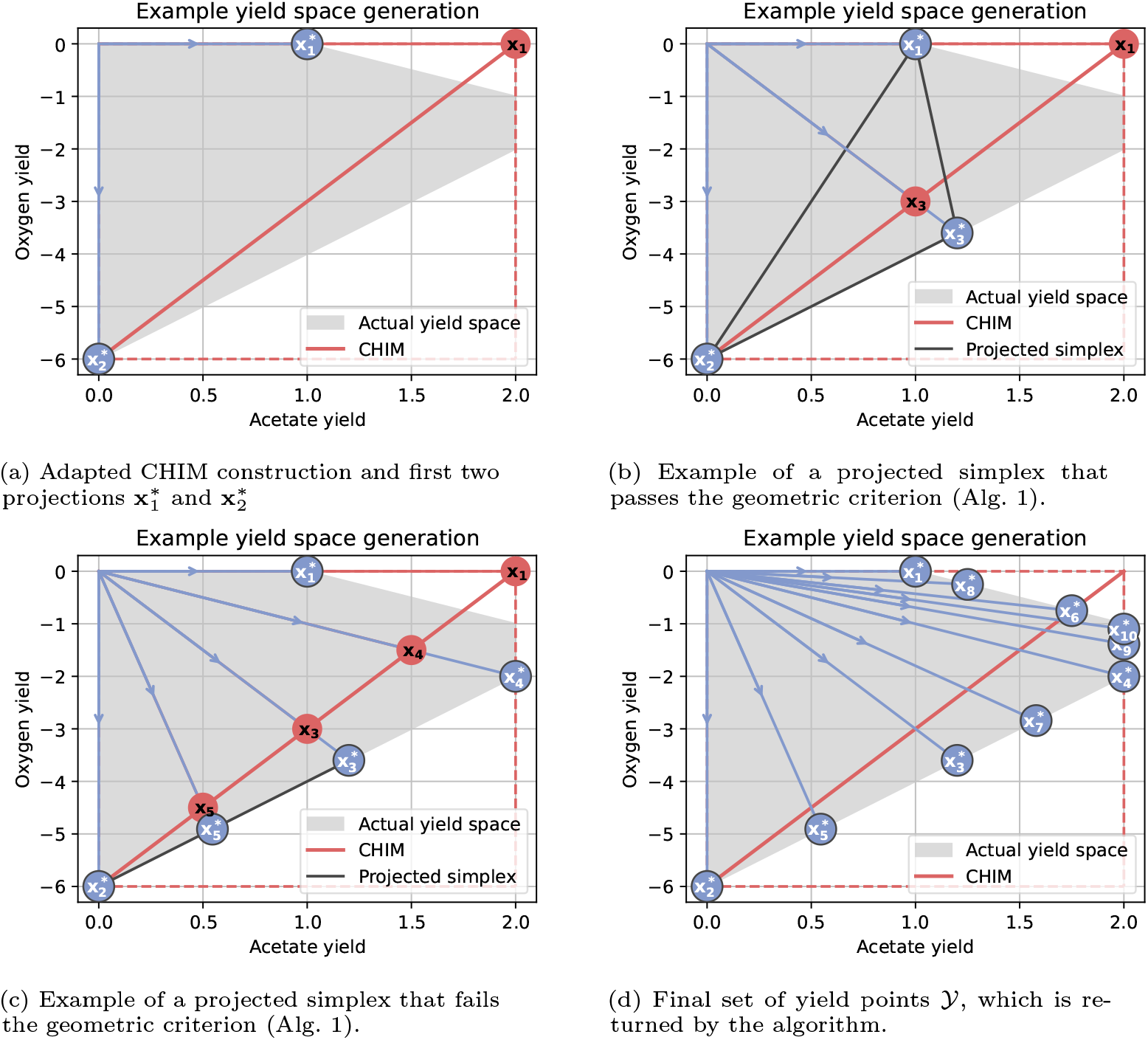
Visualization of different stages of Algorithm 2. Subfigure A shows the resulting adapted CHIM after steps 1-4, which is used to initialize the set of simplices *S*. The first subdivision of the simplices leads to the point **x**_3_ and its projection 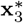 (subfigure B). Since the original projected simplex combined with the new projected point 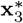 has non-zero area, the original simplex Δ_1_ = {**x**_1_, **x**_2_} is subdivided into two new simplices Δ_2_ = {**x**_1_, **x**_3_} and Δ_3_ = {**x**_2_, **x**_3_} . Subfigure C shows an example of a projected simplex which has a area of zero, which means no further subdivision is needed. Finally, once the set 𝒮 is empty, the final set of yield point 𝒴 is obtained (subfigure D). It is important to note that for higher dimensional spaces, the projected simplex would need to have non-zero (hyper-)volume.

### 2.3. Case study 1

The first case study focuses on the algorithms performance versus existing methods of generating yield spaces. For that reason, the medium-scale e_coli_core metabolic network for *E. coli* was chosen [16], allowing the generation of yield space through extreme rays. Genome-scale metabolic networks would not allow these types of methods due to the combinatorial explosion of candidates. Two yield spaces will be investigated, the first one being a 2 dimensional space regarding the biomass and acetate yield, the second one being a 4 dimensional space which also includes formate and oxygen yield. The reference substrate for all yield spaces will be glucose. The upper and lower bounds are all kept to their default for the e_coli_core network, only the glucose uptake flux was limited between *−*10 and *−*0.01, preventing zero uptake solutions. Oxygen yield is shown as negative as is standard for co-substrates, analogous to the approach for co-substrates in Song & Ramkrishna [23].

#### 2.3.1 . Compared methods

The methods compared in the first case study include the algorithm described in this work, a recent direct generation method called the opt-yield-FBA approach [8] and a recent indirect yield space generation using Extreme Pathways generated through the CBA method [14]. All methods are controlled by either the number of points sampled or by changing a tolerance such as *ϵ* (Alg. 1) which indirectly affects the yield points generated. All methods will be compared by their computation time and their volume per yield point generated.

### 2.4. Case study 2

The second case study aims to showcase the applicability of the novel algorithm described in this work to genome-scale metabolic networks and their analysis. The iML1515 network was chosen for the analysis using an *in-silico* dataset for *E*.*coli* growing in a bioreactor with limited oxygen.

#### 2.4.1. Data generation

The *in-silico* dataset was generated using a dFBA simulator based on the same genome-scale network. The simulator is based on an oxygen-limited batch bioreactor. Oxygen enters the reactor from the outside gas phase to the liquid based on te mass transfer coefficient *k*_*L*_*a* = 7.5 *hr*^*−*1^ [9]. The initial conditions are set to [0.01 *g/L*, 111 *mmol/L*, 0.21 *mmol/L*, 0.4 *mmol/L*, 0 *mmol/L*, 0 *mmol/L*] respectively. The dynamics are then as follows:

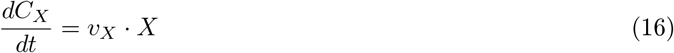

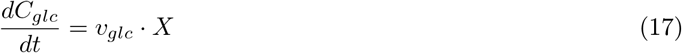

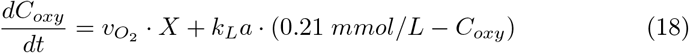

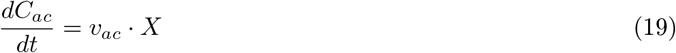

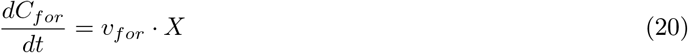

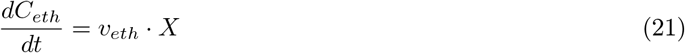

Where *C*_*X*_, *C*_*glc*_, 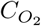, *dC*_*ac*_, *C*_*for*_, *C*_*eth*_ stand for the concentration of biomass, glucose, oxygen, acetate, formate and ethanol respectively. The unit for biomass is in *gDW/L*, while all others are in *mmol/L*, as is standard for most metabolic networks. To get the flux values, the upper limit of glucose uptake is first calculated based on a Monod kinetic with maximal rate of *−*10 *mmol/gDW/L* and a *k*_*max*_ of 5.55 [22]. Therefore, the glucose flux is bounded as:

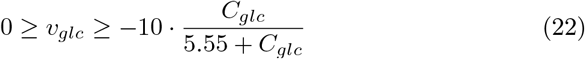

With this uptake kinetic, FBA simulation is then performed to obtain the fluxes to integrate the system of ODEs and get the state trajectories. Integration is performed until there is not enough glucose for growth and the concentration values receive 5% log-normal distributed artificial noise to simulate measurement noise. Measurement frequency is set at twice per hour.

#### 2.4.2. Exchange flux estimation

To convert the measured behaviour of cells to the yield space plots, an estimation of the fluxes has to be made based on the noisy concentration dataset. This is done by using the B-spline estimation routine as described in Vercammen et al. [25] as it provides robust flux estimation of noisy concentration datasets. The method is set to estimating the exchange rates instead of the free fluxes of a metabolic network, replacing the stoichiometric matrix with an identity matrix.

#### 2.4.3. Yield point selection

With the estimated fluxes, the yield points which are most suited to reconstruct the observed behaviour can be selected. This is done using an approach analogous to Mores et al. [13], which employs different optimization problems, estimating the weights of each yield point needed to reconstruct the observed flux. The objective for this optimization problem is then:

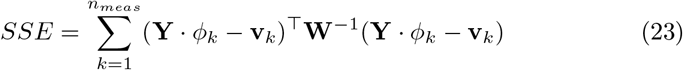

Where the corresponding flux vectors for the current yield points are combined into the matrix **Y** with shape *n*_*states*_ *× n*_*points*_, **v**_*k*_ is the estimated flux at the *k*th measurement, and *ϕ* is a vector of the macro fluxes or weights of each yield point flux vector. **W** is a diagonal matrix of shape *n*_*states*_ *× n*_*states*_ containing the maximal estimated flux for each state.

An initial sweep of the yield points is done, every time disabling just one yield point and checking how it affects the *SSE* value. If there is no significant difference, then the yield point remains disabled. After this sweep, the current set of yield points is then changed to only include the active yield points. Afterwards, the least important or worst-performer is removed from the set of yield points until a desired number of final points is achieved as defined by the user, analogous to Mores et al. [13].

#### 2.4.4. Macro-flux estimation

With a final selection of yield points, the concentration dataset can be reconstructed by using the same B-spline based approach from Section 2.4.2. Instead of estimating the exchange fluxes, the macro fluxes through each yield point is estimated by changing the identity matrix with the **Y**_*chosen*_ matrix. The fit with regards to the noisy concentration dataset is then finally evaluated, verifying whether the selection of yield points is still representative regarding the process.

## 3. Results & Discussion

### 3.1. Case study 1

The e_coli_core network [16] is analyzed in the first case study for the comparison of the new algorithm with existing, state-of-the-art methods. The Opt-yield-FBA approach [8] serves as the benchmark for direct yield space generation methods. The number of points generated is varied to evaluate its efficiency per point in terms of (hyper-)volume covered. This can be done directly through varying the number of equal-distance samples between the minimal and maximal theoretical value of the biomass yield. The total number of points generated for a 2D yield space are then 2 *· n*_*samples*_, as the non-biomass yield is both maximized and minimized while the biomass yield is fixed at the sampled value.

Extreme ray-based approaches are represented with partial EP generator [14]. The number of points generated can be controlled by the filter value *K*, which limits the number of intermediate candidates. However, the number of EPs returned is affected significantly by its stochastic nature. With the EPs, the yield space can be generated by evaluating the yields of interest with regards to the reference substrate for each EP. Afterwards taking the convex hull leads to the EP-based yield space.

The recursive approach described in this work can be scaled by changing the volume cut-off value *ϵ* in the geometric stopping criterion (Alg. 1). This will affect the depth level of recursion and therefore affect the number of yield points generated. Compared to the Opt-yield-FBA approach, the computational effort per point will be similar as it also relies on iterative LP solves, although the recursive approach has a more complex LP.

To evaluate the percentage of yield space coverage, the recursive and Optyield-FBA approaches are compared to the total yield space calculated by the EFVs [24]. In contrast, the EPs subsets generated by partial enumeration are compared to the maximal possible yield space using EPs by running the algorithm with *K* disabled. This is done as EPs generally lead to a slightly larger yield space as they do not have to adhere to inhomogenous constraints such as ATP maintenance requirements, while the direct approach and EFVs do [8, 7].

#### 3.1.1. 2D yield space

The first yield space investigated is the 2-dimensional biomass and formate yield space, with glucose as the reference substrate. All three methods are evaluated for different number of points and their performance is shown in Figure 5.

These low-dimensional kind of yield spaces are what Opt-yield-FBA was designed for, showing great performance in contrast to the indirect yield space generation of the EPs. The stochastic nature of the partial generation of EPs is very visible here, showing much less consistent coverage per point. Comparing the direct yields space generation approaches, it can be seen that the recursive approach is a slight improvement. This is expected, as the recursive approach has the ability to focus on regions of interest, increasing efficiency per point generated. This phenomenon is shown visually in Figure 4, with a limit of 10 on points generated for the direct methods. However, due to the low dimensionality of the yield space, the advantage of the recursive approach is not a significant improvement over opt-yield FBA.

**Figure 4.**
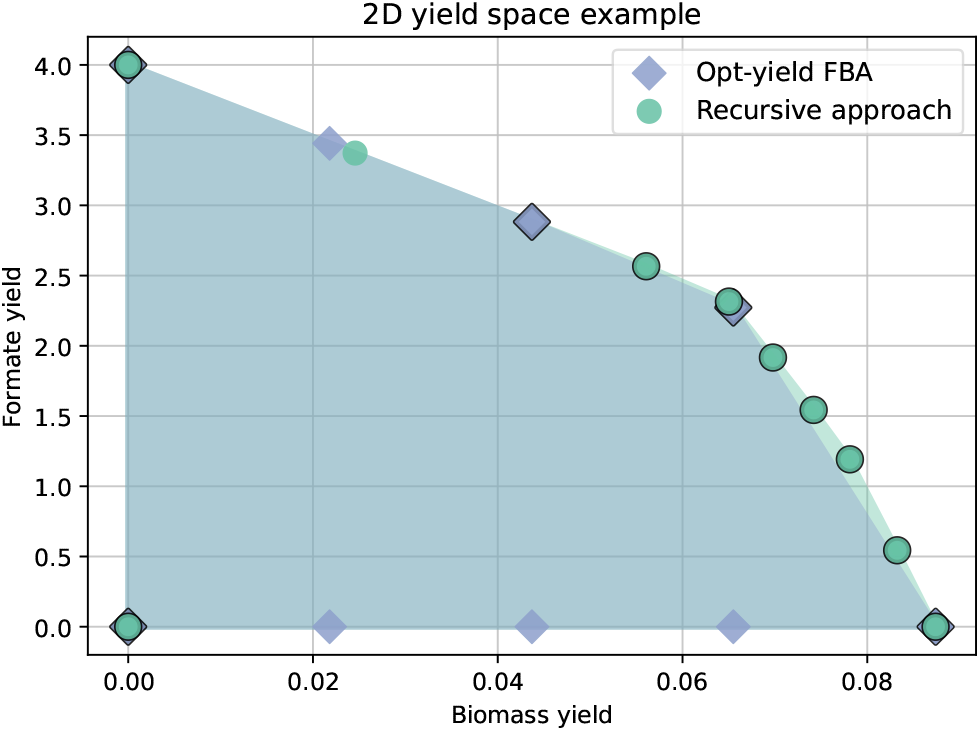
The 2 dimensional yield spaces created in case study 1. For easy comparison, both direct methods are allowed to generate 10 points to determine the yield space. Points which end up as part of the convex hull are given a black outline. For this yield space, opt-yield FBA generates significantly more insignificant points which do not contribute to the convex hull. The recursive approach finds a larger yield space and only generates 1 redundant point, after which it stops exploring the corresponding area.

#### 3.1.2. 4D yield space

To verify the applicability to high-dimensions for the recursive approach, a 4-dimensional yield space is also investigated. The yields of interest from the 2-dimensional space are extended with the acetate and oxygen yields. Similar to Song & Ramkrishna [23], oxygen is considered a co-substrate and is therefore added as a negative yield. A more negative yield then indicates that more oxygen is co-consumed to metabolize glucose.

To get a higher-dimensional coverage for Opt-yield-FBA, many 2-dimensional yield spaces are constructed and combined. Each yield of interest besides biomass will get its own yield space, with biomass yield as one axis and one yield of interest as the other. For all these points, the higher-dimensional location is determined by evaluating the flux vector and calculating the respective yields. The hyper-volume is then calculated by taking the convex hull in the high-dimensional space. This follows the approach outlined by Luo et al. [8]. For the EP and recursive approaches, no special treatment is needed as they are high-dimensional by default.

Figure 6 shows the coverage per point for all three methods for this 4-dimensional yield space. Opt-yield-FBA seems to scale relatively well, but flattens out at around 90% coverage. Afterwards, no increase of points generated will lead to the final 10% of hyper-volume. This is a drawback of the method and is also mentioned in their paper [8]. The recursive approach does reach the 100% of coverage and scales well in terms of coverage per point. Compared to the EP approach, it needs significantly less points to get full coverage. However, due to the need of solving LPs, the computational effort per point is higher for the recursive approach. Hence, the computation times are also taken into account in Figure 7.

**Figure 5.**
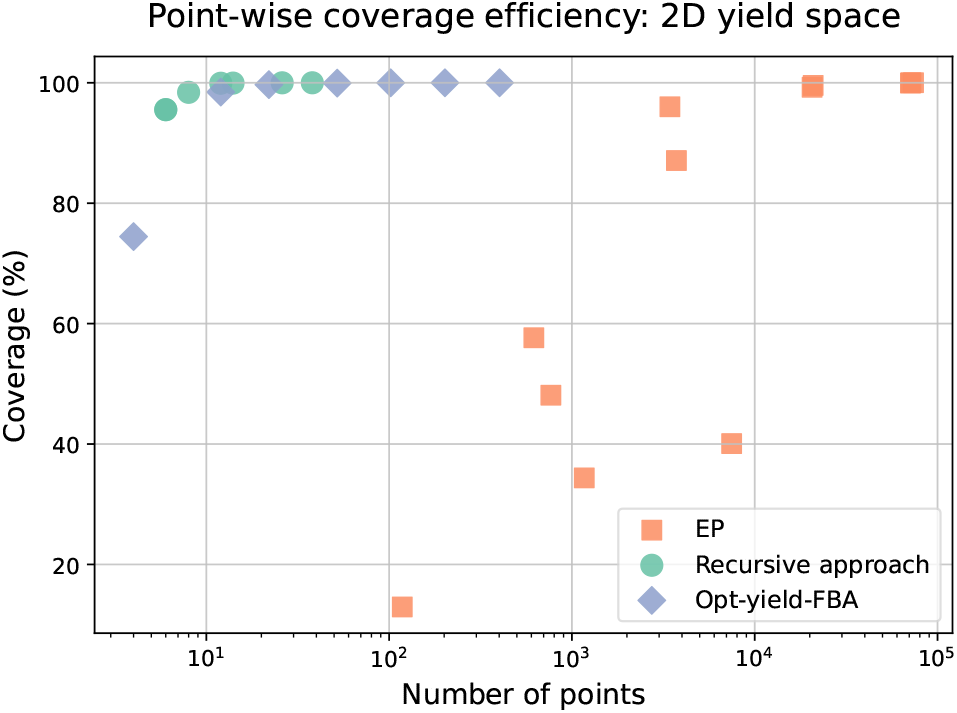
Efficiency per point in terms of yield space coverage for the e_coli_core network. The yield space has 2 dimensions, the formate yield and biomass yield. The direct generation methods scale predictably with the number of points, both reaching near perfect coverage with little amount of points. The EP approach is much more stochastic, but also leads to full coverage but requires many more points. However, it has much less computation time per point.

**Figure 6.**
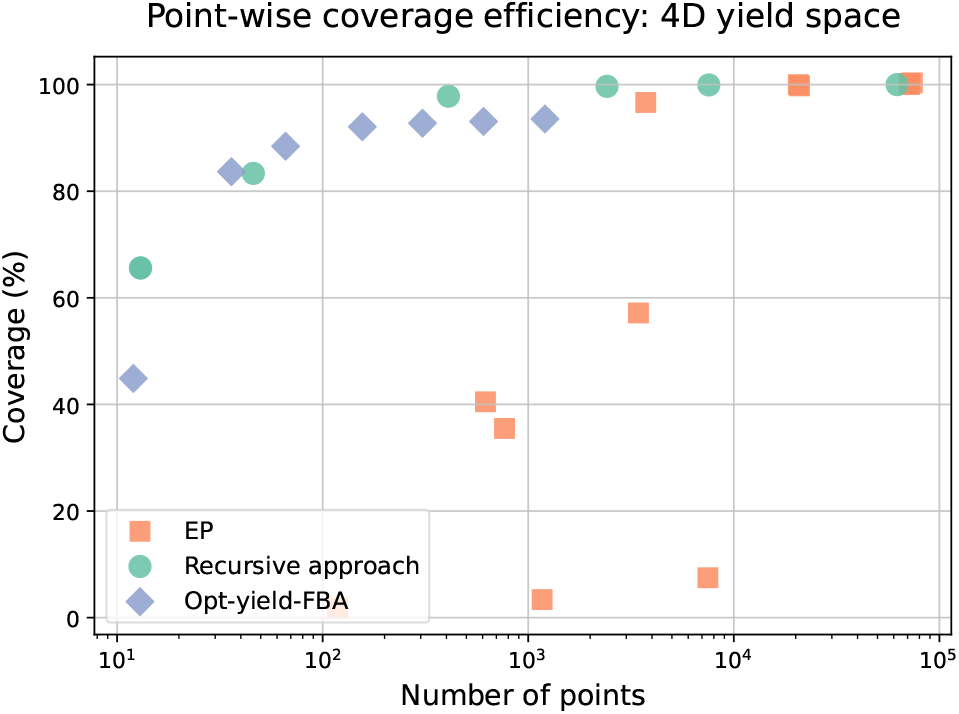
Efficiency per point in terms of yield space coverage for the e_coli_core network. A 4-dimensional yield space is now generated with the different methods. The direct generation methods again show a stable evolution of coverage with more points generated, while the EP method is more stochastic and needs significantly more points. In this higher dimensional yield space, the advantage in efficiency with our method is now clearer. When compared to the correct hyper-volume from EFV analysis, our method is capable of fully exploring. Opt-yield-FBA can only reach 93% coverage when compared to the hyper-volume of the EFVs.

**Figure 7.**
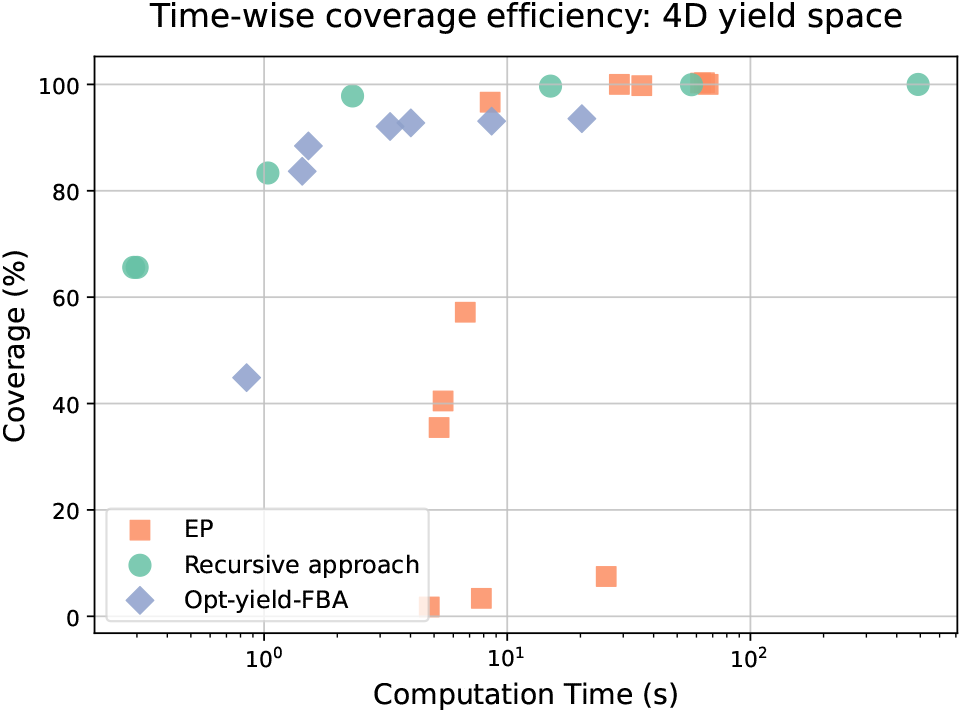
Efficiency per second in terms of yield space coverage for the e_coli_core network network.A 4-dimensional yield space is generated with the different methods. Even though the recursive approach increases the complexity of the LP, the computation time is not significantly affected. For the EP approach, the computation time per point is much lower, however, not enough to make it competitive with the recursive approach.

Considering computational effort needed for full coverage, the EP approach is much closer in performance to the recursive approach. However, if only 99% coverage or less is needed, the recursive approach needs far less time. In addition, it scales quite predictably with increasing computation time, unlike the EP approach which is very stochastic. Even for 7-10 seconds, the EP approach sometimes has less than 50% coverage while the recursive approach would give between 98 and 99%. The added complexity to the LP due to the reformulation does not seem to affect computation time per point. Both the recursive and Opt-yield-FBA approaches are built upon CobraPy [2] and are using the same solver, which should result in a fair comparison. As expected, the time per point generated for the EP approach is significantly lower, but not enough to counteract the lower efficiency in coverage per point.

### 3.2. Case study 2

Using the recursive approach for directly generating yield spaces, high-dimensional case studies can now be fully exploited even for GEMs. The iML1515 network [12], the currently most comprehensive GEM for *E. coli*, is analyzed using simulated, *in-silico* process data as described in Section 2.4. The data is generated based on the ODE system defined by Eq. 16-21, with added log-normal noise and an assumed sampling frequency of twice per hour. A 5-dimensional yield space is generated for the iML1515 model based on the state vector from the ODE system, with 4 products, one co-substrate and one reference substrate.

#### 3.2.1. Selection of yield points

Initially, the set of yield points *𝒴* resulting from Algorithm 2 contained 7745 points. After taking a convex hull and only keeping the vertices, this was reduced to 1467 points. Since 6 measurements are considered, the entire dataset should be explainable in terms of fluxes with 6 or less yield points [13, 10].

After estimating the fluxes with the B-spline-based approach, yield point selection can be done. The initial sweep, where each yield point is deactivated to check its influence on the SSE value, reduces the yield points to 156 points. The iterative worst-performer removal then allows the set to be reduced further. The evolution of the SSE from 12 to 3 points is shown in Figure 8. Based on the shape of the graph, a subset *𝒴*_*chosen*_ of 4 points offers an interesting trade-off between model complexity and accuracy for a macro-bioreaction-based model. Therefore, these 4 yield points will lie at the basis for the macro-flux estimation, potentially allowing a greatly reduced model focused on just the relevant metabolism for this specific process.

**Figure 8.**
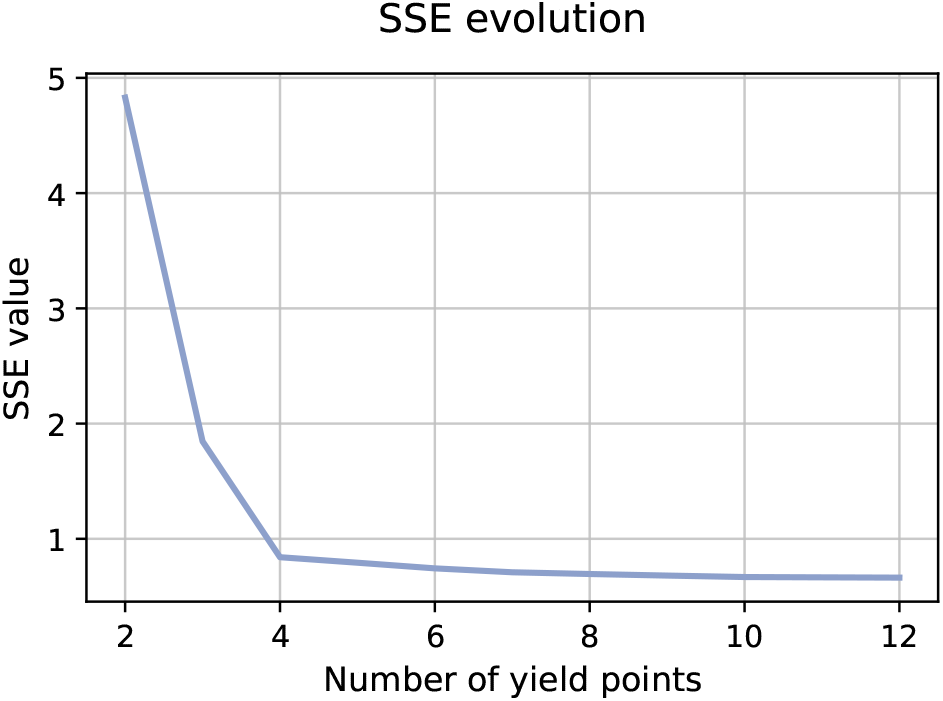
SSE values at different steps in the iterative worst-performer selection step.

#### 3.2.2. Macro-flux model

Considering the macro-conversion of each yield point in *𝒴*_*chosen*_, the matrix **Y**_*chosen*_ ∈ *n*_*states*_ *× n*_*chosen*_ is easily constructed. As described in Section 2.4.4, the macro-fluxes can be estimated using B-splines by exchanging the identity matrix used for exchange flux estimation with the matrix **Y**_*chosen*_. The resulting fit on the simulated concentration data can be seen in Figure 9. Overall, the fit is accurate, while not overfitting on the added log-normal noise, indicating that the selected yield points can fully explain the process.

**Figure 9.**
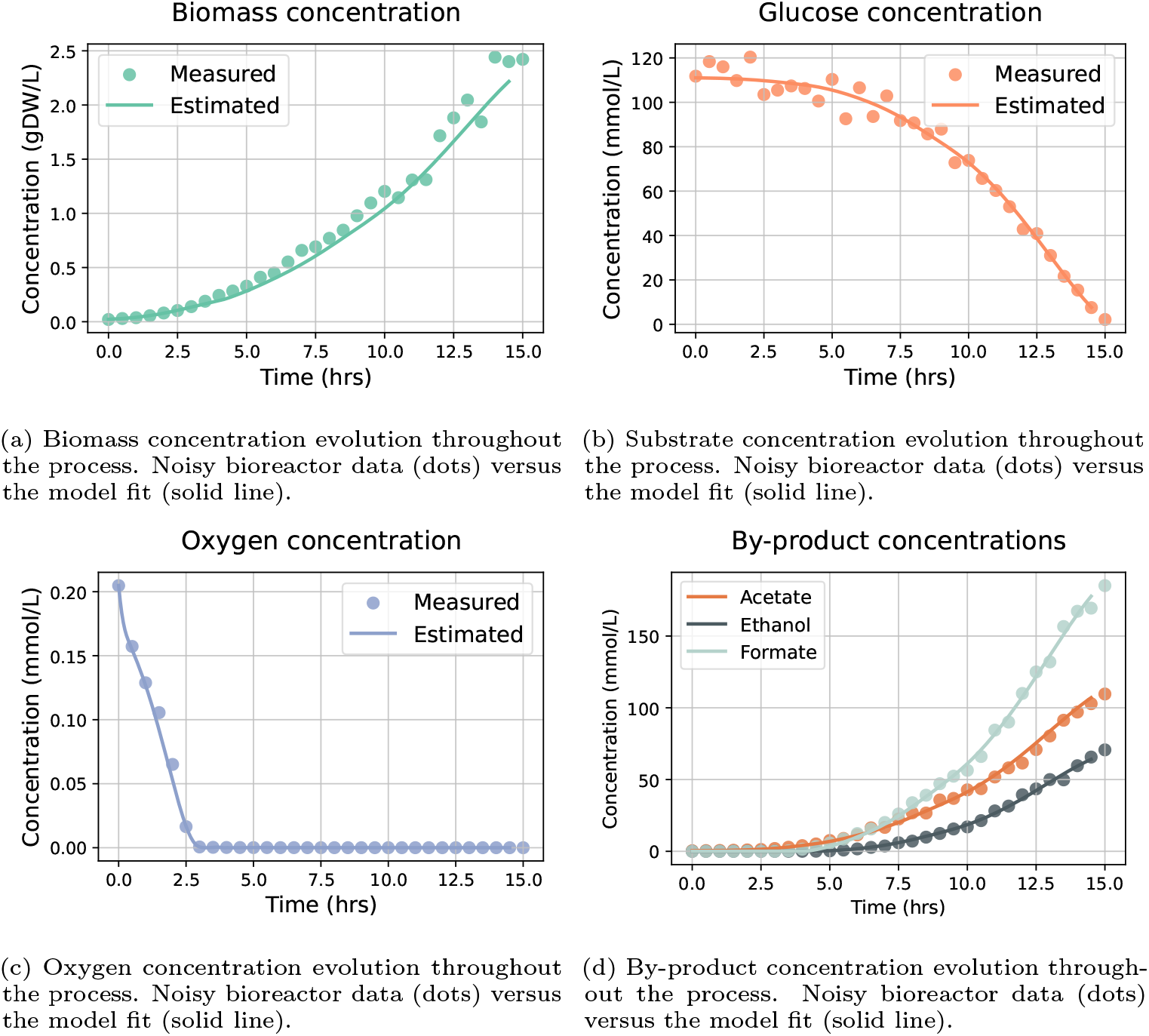
Model fit (solid line) for the macro-flux model based on the 4 selected yield points.

The macro-flux through each yield point can easily be retrieved from the B-spline estimation routine and is shown in Figure 10. For a fair comparison, each macro-flux is normalized with its carbon uptake. Considering the oxygen levels, the initial concentration depletes quickly. Afterwards, it will slowly evolve from micro-aerobic to anaerobic metabolism even though it is not visible on the concentration level. The oxygen transfer as described in Equation 18 is taken up instantly and cannot accumulate visibly on the concentration level. Anaerobic metabolism will be enabled once a certain level of biomass is achieved and the transferred oxygen has to be shared among many cells. This switch in metabolism is visible in the macro-fluxes, with the first and second yield point seemingly tied to purely aerobic metabolism, the third yield point to the micro-aerobic metabolism, and the fourth linked to the anaerobic metabolism.

**Figure 10.**
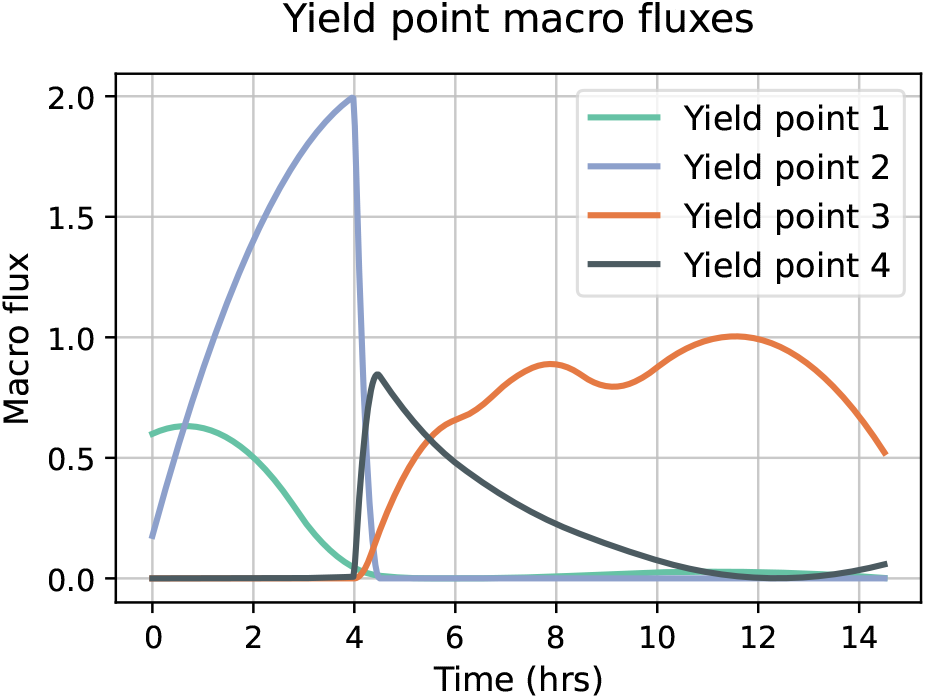
Macro-flux evolution for each yield point throughout the process, linked to the concentration evolution in Figure 9.

## 4. Conclusion

With the rise in complexity of metabolic networks, classical analysis techniques such as elementary flux mode analysis (EFM) have become less applicable. For genome-scale metabolic networks, the full set of EFMs or other types of extreme rays [7] can not be computed. Sampling within the set of extreme rays is possible [14, 17], but cannot guarantee full coverage of relevant metabolic functions, which is necessary in many applications. Yield analysis is a technique for EFM selection, allowing significant reduction of EFM candidates while keeping relevant aspects of the metabolism [13, 23]. This is done in many applications such as creating reduced models for bioprocesses. One way of avoiding the computational burden of GEMs is by directly generating the yield spaces, which removes the need of calculating the EFMs first [8]. However, the full characterization of the space is currently limited to 2-dimensional or 3-dimensional yield spaces.

In this work, a novel algorithm is presented for the direct generation of highdimensional yield spaces. By employing ideas from multi-objective optimization [15, 3], a recursive exploration method specifically catered to the properties of yield spaces is defined and tested in relevant case studies. The first case study focuses on benchmarking the novel approach to existing approaches, therefore a medium-scale network is chosen such that classical extreme ray analysis is still feasible. The second case studies then serves to show the approach is computationally feasible for recent, highly detailed genome-scale metabolic networks.

### 4.1. Efficient direct generation of high-dimensional yield spaces

Based on the case studies, the novel recursive approach has shown its ability to efficiently explore yield spaces. In low dimensional yield spaces, it performs slightly better in terms of per-point efficiency and computation times when compared to both the direct generation method Opt-yield-FBA [8]. Additionally, the novel approach only stops generating points until the geometric criteria are satisfied, giving a certain guarantee of finding the corner points of the yield space.

For higher dimensions, Opt-yield-FBA is known to have limitations and cannot explore the full yield space. The novel approach does allow full exploration, reaching 100% coverage of the theoretical maximum yield space as determined by the EFVs with few points compared to the EP-based approach. The novel recursive exploration approach is considered more efficient than both Opt-yield-FBA and the EP-based approach, even for the medium-scale network.

### 4.2. Enabling metabolic network analysis techniques for the genome-scale

By directly generating the yield space through LP solving, the novel approach is computationally feasible for genome-scale metabolic networks. For these types of networks, theoretical coverage cannot be checked as the extreme ray-based approaches are computationally feasible. Therefore, the applicability is checked with biologically relevant case study. A significantly reduced model is constructed based on a high-dimensional yield space for the iML1515 model [12], which is the most complete network for *E. coli* K-12 MG1655 to date. Using an established selection procedure for EFMs and EPs [13, 10], a selection of yield points is done. The final set of yield points contained just 4 points, where each of the macro-fluxes through the associated pathway is estimated and easily interpreted. This work showcases how the novel approach to yield space generation finally allows the use of metabolic network analysis techniques for genome-scale metabolic networks.

## Acknowledgments

This work is funded by the Research Foundation Flanders (FWO) under project G0A9H26N (*µ*4C), and the European Union under grant agreement 101122224 (ALFAFUELS). Wannes Mores was supported by Research Foundation Flanders (FWO) through Strategic Basic Project 1SHG124N. Stylianos Floros was supported by Research Foundation Flanders (FWO) through Strategic Basic Project 1SACZ26N.

## Data Availability Statement

The recursive metabolic yield space explorer is available online on GitLab KU Leuven and can be installed with **pip install git+**https://gitlab.kuleuven.be/biotec-plus/yield-space-exploration.

