## Supplementary Material for "Recursive exploration of metabolic yield space"

Wannes Mores

December 2025

#### 1 Introduction

This document provides the necessary steps to reformulate the yield LP problem for metabolic networks as defined in Luo et al. (2023) to a NBI-like LP which allows the maximization along a specific direction in the yield space. These newly defined LP's are the basis for the recursive exploration method of metabolic yield spaces.

#### 2 Yield optimization algorithm

Original yield LP problem:

$$\max \quad (v_p - Y_{temp} \cdot v_s) = 0 \tag{1}$$

$$\text{s.t.} \quad \mathbf{S} \cdot \mathbf{v} = 0 \tag{2}$$

$$\mathbf{v}_{lb} \leq \mathbf{v} \leq \mathbf{v}_{ub} \tag{3}$$

$$v_p \geq 0 \tag{4}$$

$$v_s > 0 \tag{5}$$

Where $v_p$ is the product flux, $v_s$ is the substrate flux, and $Y_{temp}$ is the current guess of the product yield, updated after every LP solve. The iterative solving stops after $Y_{temp}$ has a constant value and the objective value becomes 0.

Due to this reformulation, the fractional program created by directly using yield is avoided. This makes solving the problems significantly easier. In the original paper, only one product is considered at a time. Higher dimensional yields are explored by constraining the other while optimising one of them. To move towards NBI and more efficient multi-dimensional yield exploration, a vector of products $\mathbf{v}_p$ will be used.

### 3 Reformulation to NBI

For NBI, we change the objective to a distance metric $d$ (Das & Dennis, 1998). A constraint is then added, which involves the quasi-normal $\lambda$ and the point $P$ on the CHIM currently selected.

$$\max \quad d \tag{6}$$

$$\begin{aligned} \text{s.t.} \quad P + d \cdot \lambda &= \frac{\mathbf{v}_p}{v_s} && (7)\\ \mathbf{S} \cdot \mathbf{v} &= 0 && (8)\\ \mathbf{v}_{lb} \leq \mathbf{v} &\leq \mathbf{v}_{ub} && (9)\\ \mathbf{v_p} &\geq 0 && (10)\\ v_s &> 0 && (11) \end{aligned}$$

However, now the nolinearity is present in the constraints. By applying the same trick Luo et al. (2023) used to Eq. 7, we can try to reformulate it into a linear problem. The new constraint would then be:

$$P \cdot v_s + d \cdot \lambda \cdot v_s = \mathbf{v}_p \tag{12}$$

In this case, the multiplication of $d$ and $v_s$ makes it a bilinear equation, which is a problem for LP solvers. Substituting $d \cdot v_s$ with a new variable $e$ gives an LP which is linear in the constraints:

$$\max \quad \frac{e}{v_s} \tag{13}$$

$$\begin{aligned} \text{s.t.} \quad P \cdot v_s + e \cdot \lambda &= \mathbf{v}_p && (14)\\ \mathbf{S} \cdot \mathbf{v} &= 0 && (15)\\ \mathbf{v}_{lb} \leq \mathbf{v} &\leq \mathbf{v}_{ub} && (16)\\ \mathbf{v_p} &\geq 0 && (17)\\ v_s &> 0 && (18) \end{aligned}$$

To eliminate the nonlinear objective function, we can now apply the same trick from the paper and get:

$$\max \quad (e - Y_{temp} \cdot v_s) = 0 \tag{19}$$

$$\text{s.t.} \quad P \cdot v_s + e \cdot \lambda = \mathbf{v}_p \tag{20}$$

$$\mathbf{S} \cdot \mathbf{v} = 0 \tag{21}$$

$$\mathbf{v}_{lb} \leq \mathbf{v} \leq \mathbf{v}_{ub} \tag{22}$$

$$\mathbf{v}_{\mathbf{p}} \geq 0 \tag{23}$$

$$v_s > 0 \tag{24}$$

Where $Y_{temp}$ is recalculated every iteration with $Y_{temp} = \frac{e}{v_s}$. As a result, we now have defined an LP which is solved iteratively until the objective function is 0, which means that the correct value for $Y_{temp}$ is found. Using this formulation, yield space exploration becomes much more computationally efficient.
